# Improving the Virtual Trichrome Assessment through Bridge Category Models

**DOI:** 10.1101/2021.10.30.466613

**Authors:** Joshua Levy, Carly Bobak, Nasim Azizgolshani, Xiaoying Liu, Bing Ren, Mikhail Lisovsky, Arief Suriawinata, Brock Christensen, James O’Malley, Louis Vaickus

## Abstract

Non-alcoholic steatohepatitis (NASH) is a liver disease characterized by excessive lipid accumulation and disease progression is typically assessed through inspection of a Trichrome stain for Fibrosis staging. As the public health burden of NASH worsens due to evolving lifestyle habits, pathology laboratory resources will become increasingly strained due to rising demand for specialized stains. Virtual staining processes, computational methods which can synthesize the application of chemical staining reagents, can potentially provide resource savings by obviating the need to acquire specialized stains. Virtual staining technologies are assessed by comparing virtual and real tissue stains for their realism and ability to stage. However, these assessment methods are rife with statistical mistreatment of observed phenomena that are difficult to account for. *Bridge category ratings* represent a phenomenon where a pathologist may assign two adjacent stages simultaneously, which may bias and/or reduce the power of research findings. Such stage assignments were frequently reported in a large-scale assessment of Virtual Trichrome technologies yet were unaccounted for since no statistical adjustment procedures existed. In this work, we provide an updated assessment of Virtual Trichrome technologies using *Bridge Category Models*, which account for these *bridge ratings*. We report that two of four pathologists tended to assign lower Fibrosis stages to virtually stained tissue while the other two pathologists assigned similar stages. These research findings differ when *bridge ratings* are not accounted for. While promising, these results indicate further room for algorithmic finetuning of Virtual Trichrome technologies.

## Introduction

We present to the readership an updated assessment of the clinical viability of the Virtual Trichrome technology, previously featured as an original article in Modern Pathology. The authors, Levy et. al., previously presented a large scale validation study of virtual staining technologies with applications to the digital conversion of H&E stains to trichrome stains for the study of non-alcoholic steatohepatitis^1^. The main thrust of the research was to demonstrate concordance between stages assigned to virtual stains and stages assigned to the real stains. However, this effort was further complicated by repeat measurements, through tests and re-tests of real stains and multiple rater assessment of the same case, as well as the assignment of bridge ratings, which occurred in two-thirds of the cases^2^.

Bridge ratings represent a phenomenon in which a pathologist feels uncertain about which of two adjacent stages should be assigned. Such ratings can arise in the following ways: 1) ambiguous staging criteria, necessitating the placement of additional categories; and 2) scant information or spatially variable prognostic features across the slide may lead the pathologist to hedge against a definitive prognosis (e.g., rate fibrosis as 2-3 instead of either 2 or 3 alone). Additionally, the training and expertise of the rater may factor into this decision making. From interviews with practicing pathologists, pathologists indicated the preference to hedge against the potential for more advanced stage. Nonetheless, reporting bridge ratings are uncommon in clinical research studies, are often discouraged, and are often avoided in analyses by removing them from the analysis in some fashion ^3,4^. In the original work, Levy et. al. resorted to the application of *ad hoc* procedures, reporting concordance/correlation metrics after up/down-staging bridge categories since no statistical procedures existed for their adjustment at the time of publication ^1^.

Since then, statistical procedures have been developed to account for bridge categories, dubbed the *Bridge Category Models* ^5,6^, further divided into separate models (*Expanded, Mixture, Collapsed*) which represent different data generating mechanisms. Of note, the *Mixture Bridge Category* model allows for the report of a parameter, *p*, which indicates the propensity of the pathologist for reporting two higher adjacent stages (bridge ratings; *Y* = {*j, j* + 1}) when they may have felt the features of the biopsy were indicative of the lower of the two adjacent stages (*Y* = *j*) (*1-p* would indicate the opposite effect). Simulation studies of the *Bridge Category Models* indicated that applying *ad hoc* procedures to choose a single stage from a bridge rating (either randomly, or systematically upwards/downwards) would likely lead to biased or imprecise effect estimates ^5,6^. However, if the reason why the pathologist assigned a bridge rating is known (e.g., hedging against a more severe stage), then selecting a *Bridge Category* model that is commensurate with the data generating mechanism can circumvent issues associated with the application of *ad hoc* procedures. Recently, *Bridge Category Models* have been modified to account for ratings from multiple pathologists, indicating whether certain pathologists were more or less likely to assign the lower stage of the bridge rating ^5,6^.

In this work, we reassess the viability of virtual trichrome staining of liver for fibrosis assessment using the *Bridge Category Models*. The original model design in Levy et. al. reported how well stages assigned to the virtual stains correlated with real stains after accounting for bias. In that paper, bridge ratings were removed prior to analysis through ad hoc procedures, which reported effects by considering separately the higher or lower stages of the bridge ratings. Here, we instead use *Bridge Category Models* to assess for systematic differences in staging preferences associated with the assessment of a virtual stain (i.e., do pathologists assign a higher or lower stage if the stain is virtual?). The same functional form is also used to predict factors related to the treatment of the bridge rating (e.g., do pathologists believe the lower stage of the bridge rating to be true if the stain is virtual?).

## Methods

We have included the specification of the statistical model here, which assesses the likelihood of obtaining stage *Y*_*i*_ = *j* or *Y*_*i*_ = {*j, j* + 1} (bridge rating) for scenarios in which a bridge rating is supplied (see Appendix section “Hierarchical Bridge Category Model Specification for Virtual Trichrome Assessment”):

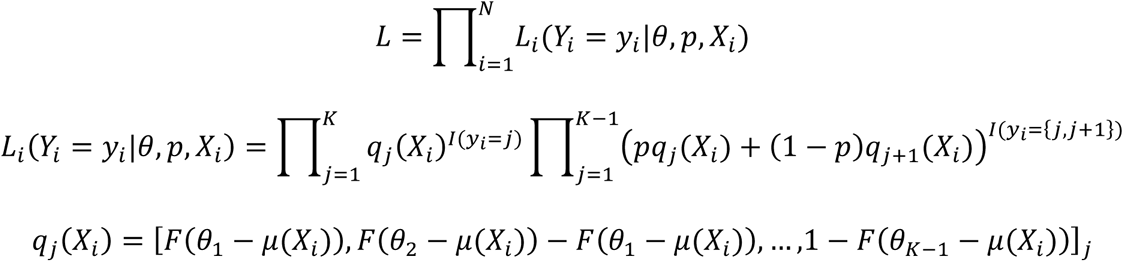

with *F*() the cumulative distribution function of the standard normal distribution,

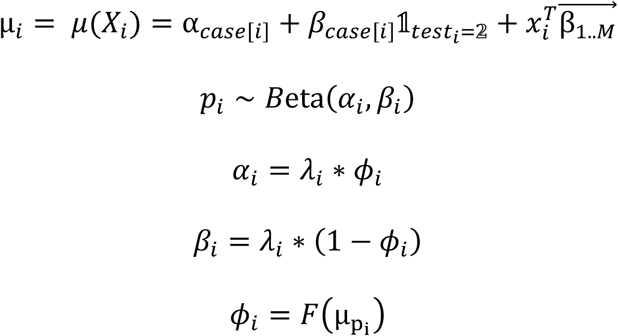

with *F*() the cumulative distribution function of the standard normal distribution,

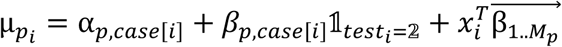

The outcome *Y* is an ordinal category that is multinomially distributed, where categorical probabilities are determined using a cumulative link model ^5,6^. A mixture parameter, *p*_*i*_, is drawn to weight the likelihood (*L*_*i*_(*Y*_*i*_ = *y*_*i*_|*θ, p, X*_*i*_)) probabilities *q*, based on whether the pathologist felt that, on average, the true rating is the lower of the two adjacent categories, with some dependence on which rater was staging and whether the stain was real/virtual 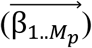. In this study, we aim to estimate the following inference targets:

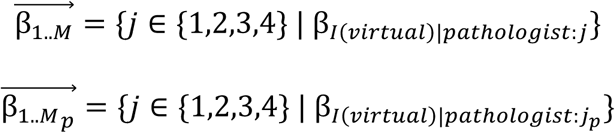

Here, *I*(*virtual*) is a flag that indicates whether the observation was from a virtual slide image. The corresponding effect estimate on this factor will report on a pathologist-specific basis: 1) whether the virtual stage was associated with systematic over/under-staging 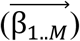; and 2) for bridge categories, the propensity for up/down staging from the bridge rating 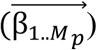, for which the mixture parameter *p* can be reported on for each pathologist and whether the tissue was real/virtual. A report of a 95% credible interval that lies outside of zero indicates a statistically significant result (systematic assignment of higher or lower stages and/or bridge ratings, both dependent on whether the stain was real or virtual). We refer to two previous works as to the formulation of *Bridge Category Models* and their hierarchical extensions and have included detail on the complete model specification featured in this work in the Supplementary Materials (section “Hierarchical Bridge Category Model Specification for Virtual Trichrome Assessment”). We fit the statistical models using hierarchical Bayesian procedures, available through R and Stan code at the following URL: https://github.com/jlevy44/VirtualTrichromeBridgeCategoryAssessment ^7,8^.

## Results

Taking into account bridge category ratings, we report that two out of the four pathologists (pathologists one and four) tended to report lower stages when confronted with a virtual stain (Table 1; Supplementary Figure 1A). Additionally, all four pathologists had a different propensity for up/down-staging bridge categories (Figure 1; Supplementary Figures 1B; Supplementary Table 1). However, for three out of four pathologists, these propensities were different depending on whether the stain was real or virtual. For instance, pathologist one tended to assign a bridge rating (two adjacent stages) while believing features were indicative of the lower of the two adjacent stages (e.g., slide showed features of 3 but staged it 3-4 anyway), regardless of whether the stain was real or virtual. Meanwhile, pathologists two and three tended to assign the lower of two stages to virtual stains while having a higher stage in mind, as compared to practices for real stains, and pathologist 4 demonstrated an opposite effect. Using the *ad hoc* procedures (upstaging, downstaging or random selection from bridge ratings), significant divergences/differences are reported after the loss of information (Supplementary Tables 2-5).

**Table 1:** Posterior effect estimates using *Mixture Bridge Category Model* for estimating whether stages assigned to virtual stains were different from real stains; Virtual vs real column indicates whether posterior credible interval indicates that either stages assigned to virtual were less than real stains (first 4 rows) or whether bridge categories assigned virtual stains were thought to the lower of two stages (last 4 rows)

|  |  | Mean | 95%CI-Low | 95%CI-High | 95%Width | Virtual vs Real |
| --- | --- | --- | --- | --- | --- | --- |
| **Stages Assigned to Virtual Stains vs. Real Stains** | Pathologist 1 | -1 | -1.17 | -0.82 | 0.35 | Less |
|  | Pathologist 2 | -0.11 | -0.36 | 0.11 | 0.47 |  |
|  | Pathologist 3 | 0 | -0.22 | 0.2 | 0.43 |  |
|  | Pathologist 4 | -1.68 | -1.88 | -1.49 | 0.4 | Less |
| **Propensity Up/Down Staging Bridge Categories** | Pathologist 1 | 0.06 | -1.23 | 1.32 | 2.55 | Down-Stage |
|  | Pathologist 2 | -1.16 | -2.21 | 0.06 | 2.27 |  |
|  | Pathologist 3 | -1.12 | -2.24 | -0.28 | 1.97 | Up-Stage |
|  | Pathologist 4 | 1.33 | 0.5 | 2.14 | 1.63 | Down-Stage |

**Figure 1:**
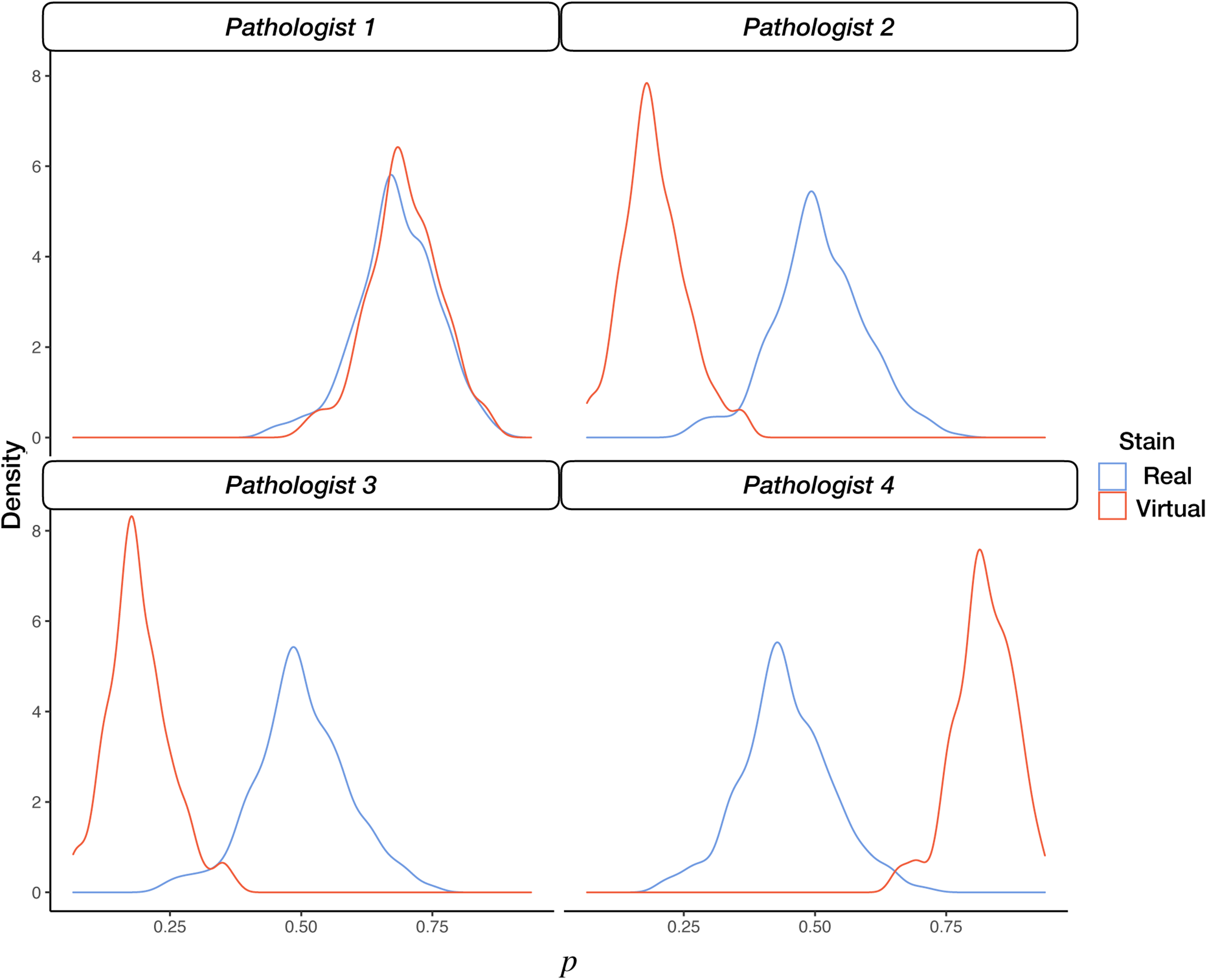
Visualization of posterior distribution (faceted density plot) of up/downstaging propensity for each pathologist and whether slide was real or virtual. When mean of distribution is closer to 1, tendency is to downstage or consider lower of two bridge categories, and when mean is closer to 0, tendency is to upstage or consider higher of two bridge categories

## Discussion

It is an interesting coincidence that pathologists who showed the least divergence between stages assigned to virtual stains versus that of real stains (pathologists two and three) also tended to assign the lower of two stages when reporting a bridge category rating (e.g., on average across all assigned bridge ratings they recorded stage 2-3 when they really meant to assign stage 3). While this new analysis further supports that virtual trichrome staining is a viable intermediate diagnostic decision aid, there is clearly further room for improvement. Regardless, the application of the *Bridge Category Model* to the assessment of Virtual Staining technologies may serve as a guide for other biomedical researchers seeking to validate their technologies using best statistical practices ^2,9^. For instance, the *Bridge Category Model* presents pathologist specific estimates of the tendency to up/downstage bridge categories, which provide useful information for finetuning staging / prognostication / drug effect technologies and understanding clinical decision making (as well as suggesting a potential new avenue for training and maintenance of certification). We believe that the Bridge Category Model approach is superior to *ad hoc* adjustments in that all available information is utilized rather than arbitrarily truncated.

Our research findings from this correspondence are consistent with reports from the Turing Test analysis (a visual quality assessment, e.g., do virtual images appear real? ^10^) in Levy et. al., where the same two pathologists who were unable to tell the difference between the real and virtual tissue also did not systematically alter staging practices for real versus virtual stains ^1^. However, utilizing the Turing test for medical images is problematic and opportunities exist to improve its clinical utility. For instance, the Turing Test does not take into account repeated measurements and multiple raters. Existing Turing Test measures are potentially misaligned with the clinical viability of the technology, as they can be gamed by presenting highly magnified images, which make it difficult to spot differences between real and virtual stains. This virtual tissue visual quality assessment (i.e., do virtual images appear real?) can potentially benefit through integration with clinical viability assessment methods (i.e., does real/virtual tissue stage similarly?). In the future, we plan to incorporate hierarchical Bayesian *Bridge Category Models* with the Turing Test method as a strategy to suggest the degree to which the quality of the tissue staining drives differences in staging patterns for real and virtual tissue, and at what scale this occurs.

## Appendix

### Hierarchical Bridge Category Model Specification for Virtual Trichrome Assessment

We refer to two prior works for the specification of the Bayesian modeling, bridge category models and their hierarchical extensions. Here, we discuss the formulation for a hierarchical bridge category model to assess for differences in staging practices due to virtual staining.

The bridge category models presented here assume that only one pathologist is staging the biopsies and that there are no repeat biopsies. However, this assumption is unrealistic given standard clinical practice and the need to estimate the reliability / validity of the stages measured. As such, hierarchical modeling can account for variation in the stages assigned by different pathologists and within specific cases. Failing to account for repeated measurements may also render the study conclusions invalid, imprecise or significantly biased. Hierarchical Bridge Category Models assume are able to estimate different values of *p* for different pathologists and cases, adding either random or fixed effects for the estimation of *p*. This essentially means that pathologists may have different up or downstaging practices when assigning bridge categories. Hierarchical *Mixture* bridge category models for this work are specified as follows:

The likelihood function for the entire sample of observations is given by:

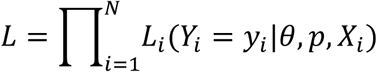

with the contribution from the ith observation given by:

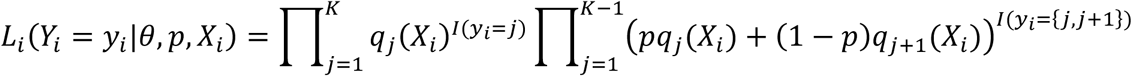

where:

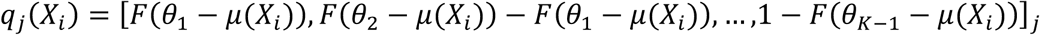

with *F*() the cumulative distribution function of the standard normal distribution,

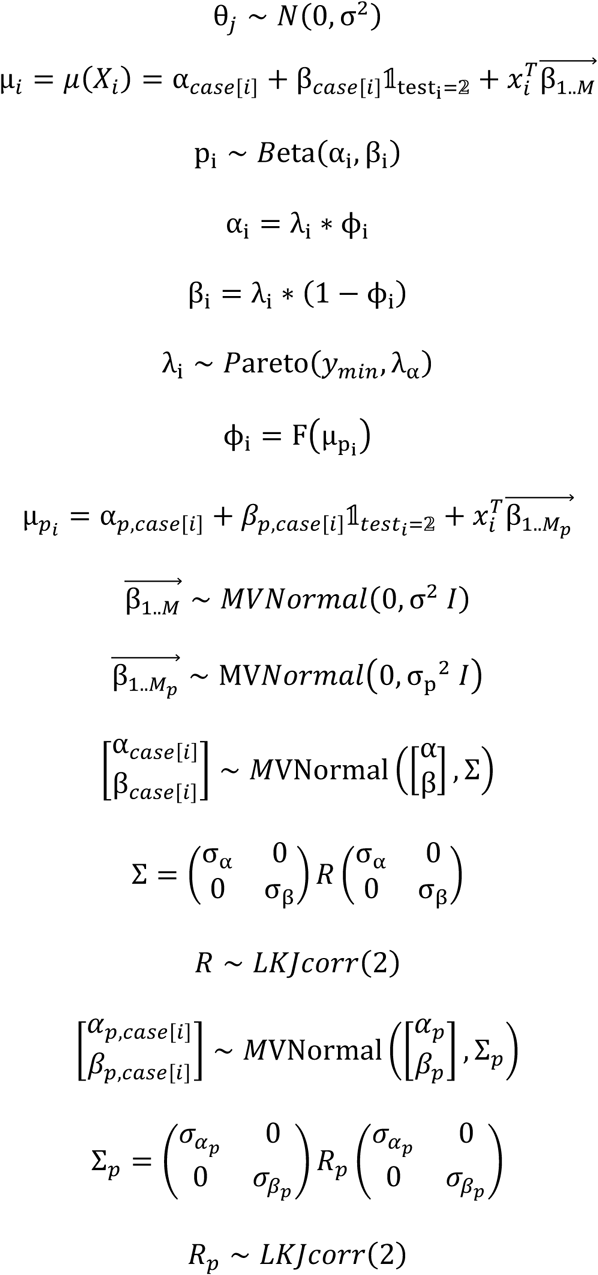

In this Bayesian model, covariate information and case level random intercepts/slopes are used to inform the estimation of parameter *p*. The cumulative distribution function *F* is used to transform the conditional mean for *p*, µ_p_, to a value between 0 and 1. Random intercepts (*α*_*case* [*i*]_, *α*_*case*[*i*]_) account for variation between cases, while random slopes (*β*_*case*[*i*]_, *β*_*case* [*i*]_) account for changes in assignments over time on a case-by-case basis, where these changes may be correlated with the random intercepts by an amount quantified by the off-diagonal element of *R* or *R*_*p*_.

For the assessment of the Virtual Trichrome technology, we wanted to demonstrate whether pathologists stage virtual tissue differently than real tissue. The main predictors of Fibrosis stage in our ordinal regression model 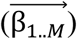 were whether the tissue stain was real or virtual (β_*I*(*virtual*)_), and which pathologist staged the tissue (β_Pathologist:2…4_). If β_*I*(*virtual*)_ is found to be less than zero, this would indicate that virtual stains were assigned lower stages than real stains. For the pathologists, staging practices from *pathologists 2 through 4* are compared to *pathologist 1*; when β_Pathologist:2…4_ is found to be greater than zero, this would indicate that the other pathologists assigned higher stages than pathologist 1. We also considered the scenario where a pathologist may be more or less likely to assign a higher or lower stage based on whether the stain was virtual by generating some interaction terms, β_*I*(*virtual*)∗Pathologist:2…4_. Thus, for each pathologist, the final estimate for each pathologist as to whether stages assigned to virtual stains were different from real stains is given by β_*I*(*virtual*)_ for pathologist 1 and β_*I*(*virtual*)_ + β_*I*(*virtual*)∗Pathologist:2…4_ for pathologists 2 through 4. The latter term is equivalent to the effect of virtual staining, conditional on the pathologist, where the reference level is with respect to pathologist 1 (β_*I*(*virtual*)∗Pathologist:1_ does not exist):

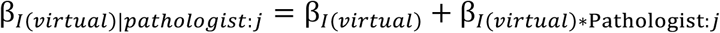

Our main targets of inference for this study were β_*I*(*virtual*)_ and β_*I*(*virtual*)|*Pathologist*:_*j*_*p*_ to assess potential differences in stage assignment, attributable to Virtual Trichrome:

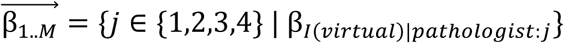

We separately wanted to assess whether, for bridge ratings, the virtual staining influenced up/downstaging practices (estimation of *p*). The estimation of the effect is similar to that estimated for impact of stain on staging, but now the estimated effects / parameters reference the population parameter *p*. The main targets of inference here were β_*I*(*virtual*)*p*_ and β_*I*(*virtual*)|*pathologist*:_*j*_*p*_, where the *p* subscript has been added to ascribe these parameters for the predisposition to up/downstage for specific pathologists:

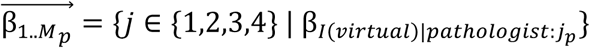

### Plots of Posterior Estimates

**Supplementary Figure 1:**
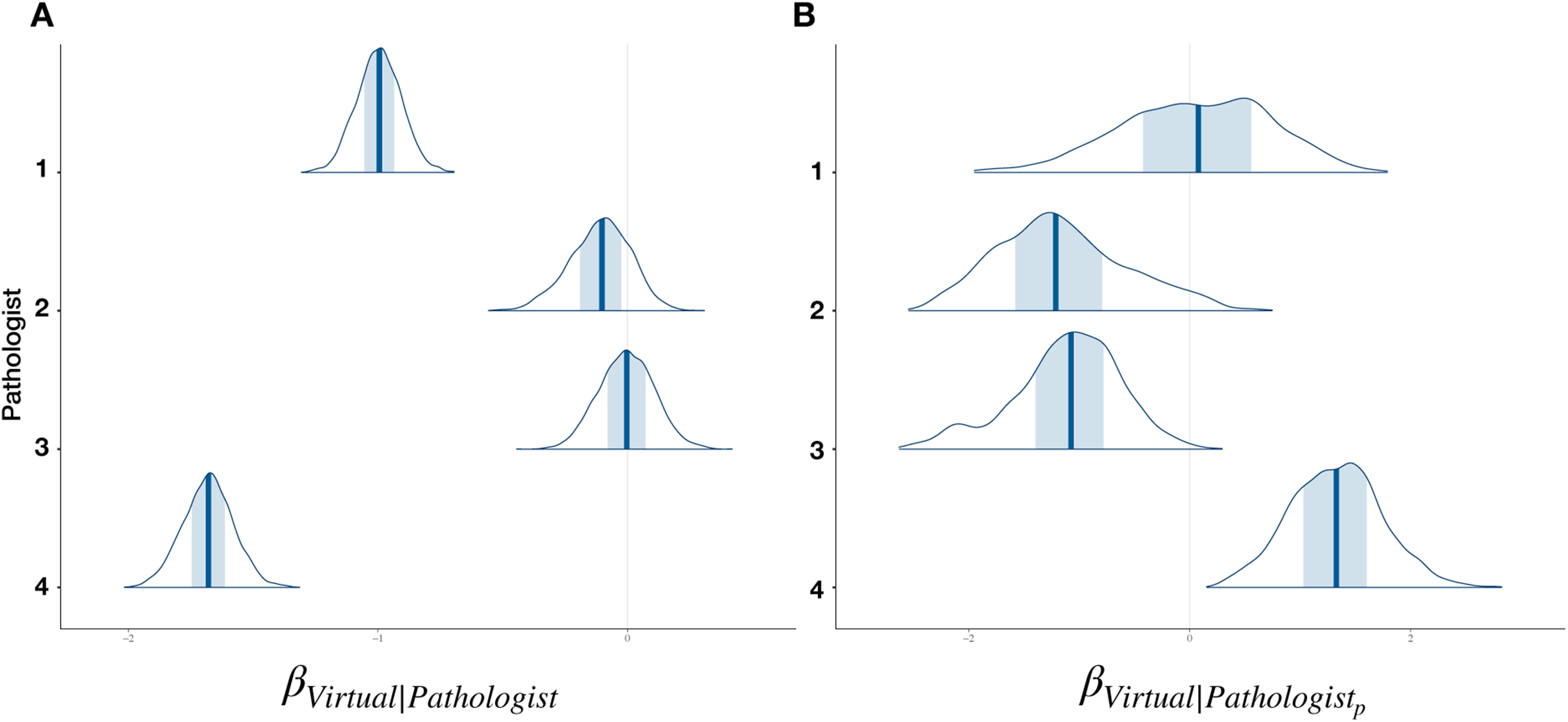
Density plots of posterior distribution of effect estimates for whether: A) Virtual stain has higher stage than real stain (greater than 0); and B) whether bridge category leaned towards down staging based on whether stain was virtual (greater than 0) or real; Posterior distribution of parameters estimated for reach pathologist by considering conditional effects via the addition of the interaction to the main effect

### Pathologist-Specific Bridge Category Up/Down Staging Propensity

**Supplementary Table 1:** Posterior estimates for mixture parameter *p* for up/down-staging propensity of bridge ratings for each pathologist

| Pathologist | Stain | Mean | 95%CI-Low | 95%CI-High | 95% Width |
| --- | --- | --- | --- | --- | --- |
| Pathologist 1 | Real | 0.68 | 0.54 | 0.86 | 0.31 |
|  | Virtual | 0.70 | 0.58 | 0.86 | 0.28 |
| Pathologist 2 | Real | 0.51 | 0.34 | 0.71 | 0.37 |
|  | Virtual | 0.19 | 0.07 | 0.31 | 0.24 |
| Pathologist 3 | Real | 0.50 | 0.33 | 0.70 | 0.37 |
|  | Virtual | 0.19 | 0.07 | 0.30 | 0.23 |
| Pathologist 4 | Real | 0.45 | 0.29 | 0.65 | 0.37 |
|  | Virtual | 0.82 | 0.72 | 0.94 | 0.22 |

### Results from Other Methods

**Supplementary Table 2:** Posterior effect estimates using *Up-Staged Ordinal Regression* for estimating whether stages assigned to virtual stains were different from real stains; Virtual vs real column indicates whether posterior credible interval indicates that either stages assigned to virtual were less than real stains

|  |  | Mean | 95%CI-Low | 95%CI-High | 95%Width | Virtual vs Real |
| --- | --- | --- | --- | --- | --- | --- |
| Stages Assigned to Virtual Stains vs. Real Stains | Pathologist 1 | -0.94 | -1.1 | -0.78 | 0.32 | Less |
|  | Pathologist 2 | -0.21 | -0.38 | -0.05 | 0.33 | Less |
|  | Pathologist 3 | -0.16 | -0.32 | 0 | 0.32 |  |
|  | Pathologist 4 | -1.43 | -1.61 | -1.26 | 0.35 | Less |

**Supplementary Table 3:** Posterior effect estimates using *Down-Staged Ordinal Regression* for estimating whether stages assigned to virtual stains were different from real stains; Virtual vs real column indicates whether posterior credible interval indicates that either stages assigned to virtual were less than real stains

|  |  | Mean | 95%CI-Low | 95%CI-High | 95%Width | Virtual vs Real |
| --- | --- | --- | --- | --- | --- | --- |
| Stages Assigned to Virtual Stains vs. Real Stains | Pathologist 1 | -0.95 | -1.11 | -0.79 | 0.32 | Less |
|  | Pathologist 2 | -0.35 | -0.51 | -0.19 | 0.33 | Less |
|  | Pathologist 3 | -0.19 | -0.36 | -0.03 | 0.33 | Less |
|  | Pathologist 4 | -1.45 | -1.62 | -1.28 | 0.34 | Less |

**Supplementary Table 4:** Posterior effect estimates using *Randomly-Staged Ordinal Regression* for estimating whether stages assigned to virtual stains were different from real stains; Virtual vs real column indicates whether posterior credible interval indicates that either stages assigned to virtual were less than real stains

|  |  | Mean | 95%CI-Low | 95%CI-High | 95%Width | Virtual vs Real |
| --- | --- | --- | --- | --- | --- | --- |
|  | Pathologist 1 | -0.93 | -1.08 | -0.76 | 0.33 | Less |
| **Stages Assigned to Virtual Stains vs. Real Stains** | Pathologist 2 | -0.29 | -0.44 | -0.13 | 0.31 | Less |
|  | Pathologist 3 | -0.15 | -0.31 | 0.01 | 0.32 |  |
|  | Pathologist 4 |  |  |  |  | Less |
|  |  | -1.4 | -1.57 | -1.24 | 0.34 |  |

**Supplementary Table 5:**
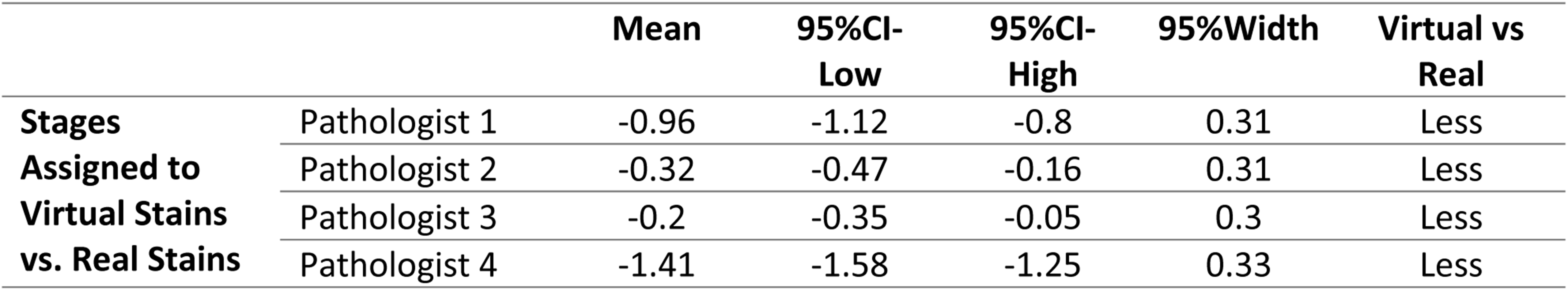
Posterior effect estimates using the E*xpanded Bridge Category Model* for estimating whether stages assigned to virtual stains were different from real stains; Virtual vs real column indicates whether posterior credible interval indicates that either stages assigned to virtual were less than real stains

